# Dendritic cell CX3CR1 and macrophages F4/80 play a central role in between gut micro biome and inflammation in Arsenic induced mice

**DOI:** 10.1101/2021.01.25.428065

**Authors:** Chiranjeevi Tikka, Ram Kumar Manthari, Ruiyan Niu, Zilong Sun, Jundong Wang

**Author notes:** **Corresponding author:** Prof. Jundong Wang, Shanxi Key Laboratory of Ecological Animal Science and Environmental Veterinary Medicine, College of Animal Science and Veterinary Medicine, Shanxi Agricultural University, Taigu-030801, Shanxi, China.

## Abstract

Microbiota plays a crucial role to protect the intestine contrary to the harmful foreign microorganisms and organize the immune system via numerous mechanisms, which include either direct or indirect environmental factors. The underlying mechanism arsenic (As) influenced immune system and regulates inflammation by altering gut microbiome in ileum remains unclear. However, chronic exposure to arsenic (at doses of 0.15 mg or 1.5 mg or 15 mg As_2_O_3_/ L in drinking water) significantly increased mRNA and protein levels of F4/80 and CX3CR1, concurrently, the increased levels of mRNA and protein IFNγ, TNFα, IL-18 and decreased levels of IL-10 were found in both 3 and 6 months exposure periods. High-throughput sequencing analysis revealed that gut microbiota at phylum; family and taxonomical levels were showed the abundance of gut microbiota. Evidentially, the ultra-structure of intestinal villi, microbes engulfed and immune cell migration were showed by the transmission electron microscopy. Chronic exposure to As influenced the inflammation by changing immune system and altered gut microbiota. In this study we conclude that chronic exposure to As breakdown the normal gut microbial community and increase the pathogenicity, the resultant risk pathogen direct contact with intestinal immune system and regulate the inflammation.

## 1. Introduction

Arsenic (As) is a one of the most dangerous chemical toxic factor in the environment; it is widely distributed in the natural weathering of the earth’s crust. Likewise, it certainly develops linked with water, soil, food and particulates in the air (Nordstrom et al., 2002). Moreover, the arsenic contaminate can occur in the region of mining and industrial arsenic application by the human being [Williams et al., 2001]. The sever exposure to high dose of As is more toxic to most cellular life forms, and prolonged low dose exposure in human being, is also linked with several disease. The community of microorganisms is called microbiome, which residing in a well-defined environment and the quantity of their physical, biological and ecological activities (Whipps et al., 1988). The human body contains trillions of microbiome, which taxonomically described as micro-eukaryotes, archea, virus, bacteria and fungi. It is ultimately linked between the abiotic environment, mcrobiome and host cells have substantial influences on human health and disease (Clemente et al., 2012). In this study we focus on alteration of gut microbiome influence inflammation via disruption of intestinal immune system in chromic exposure of arsenic.

The specific function of mucosal immune system is mainly self-growing of the universal immune system (Woof et al., 2006) and it ultimate changes occurs after bacterial migration of the gastro intestinal tract (Scheppach et al., 1994). The immune system maturation is depends on commensal microorganisms, which study to separate between commensal bacteria and pathogenic bacteria (Thaiss et al., 2016; Nakanishi et al., 2015). For the development of mucosal immune system in the intestinal epithelial cells and lymphoid cells by the active function of Toll-like receptors (TLRs). The role of TLRs to inhibits the inflammatory response and stimulates immunological tolerance to typical microbiota components. Moreover, gut microbiota control neutrophil passage, function (Owaga et al., 2015) and disturb the immune cell populations into different types of helper cells (Th), such as Th1, Th2, and Th17 or into regulatory T cells (Francino et al., 2014).

The helper cells of Th17 cells are subclass of TCD4+ cells, which secrete numerous cytokines (IL-17A, IL-17F and IL-22), with a substantial influence on immune homeostasis and inflammation (Rossi et al., 2013; Sonnenberg et al., 2011). Typically stimulate macrophages secrets a huge volume of IFN-γ, TNF-α, IL-6, and IL-12 and express inflammatory and anti-inflammatory activities. Anti-inflamatory IL-10 plays an important role in the activation of Macrophages especially in the region of intestine, in addition IL-10 plays crucial role in the deletion of inflammation, which associate with hyper responsiveness of macrophages (Rivollier et al., 2012; Hirotani et al., 2005; Takeda et al., 1999; Ueda Y et al., 2010; Kühn R; Löhler., et al., 1990; Zigmond E et al., 2014; Shouval DS., 2014). In this study we report that chronic exposure to Arsenic induced inflammation by the hyper expression of Macrophages (F4/80) and dendritic cells (CX3CR1) that stimulated inflammatory (TNF-α, sIFN-γ and IL-18) cytokines while suppressing the anti-inflammatory (IL-10) cytokines through the altered gut microbiome in the region of Ileum. However, expression of macrophages, dendritic cells and inflammatory and anti-inflammatory cytokines significantly increased and depletion levels were found at dose dependent which associated with gut microbiome alterations.

### 2. EXPERIMENTAL METHODOLOGY AND MATERIALS

### 2.1. Animal exposure and Maintenance

120 male Kunming mice of 6 weeks age were purchased from Experimental Animal Centre, Academy of Military Medical Sciences, China. All mice were allowed two weeks for environmental adaptation, maintained at 220C, 45-65% humidity, and a 12:12 hrs dark: light cycle before commencement of the experiment. During this period, mice were received only deionized water and pelleted rodent diet. After completion of the two weeks of environmental adaptation, the mice were allocated into two groups for two different exposure periods (3 months and 6 months; containing 60 mice each). 60 mice from each exposure period, were randomly divided into four groups (each group containing 15 mice) as control (received only Distilled H2O), low (0.15 mg As2O3/L), medium (1.5 mg As2O3/L) and high (15 mg As2O3/L) dose. As-containing distilled water and pelleted rodent diet were provided twice in a week. Each mouse drink about 10 mL/day on an average daily, which resulted in intake of 1.5 µg, 15 µg and 150 µg per mouse per day in low, medium and high groups respectively. These doses have been chosen according to previous studies [37] and based on environmental concentrations of As or surveillance of realistic exposure doses for humans, every group has housed in a microbe’s free cages on heat-treated hardwood bedding. All the experimental procedures have been accepted by the Animal Welfare International Ethics Committee, Shanxi Agricultural University, China.

### 2.2. Histological studies

After completion of the two exposure periods (3 months and 6 months), the mice were sacrificed and dissected entire small intestine. Then ileum part was separated and then fixed in formalin solution (10%) for 24–28 h. Next the formalin fixed samples were dehydrated in an alcohol grade series and then fixed in paraffin wax. Prepared sections (4–5µM thickness) have been stained through hematoxylin–eosin for histopathological studies.

### 2.3. Immunohistochemistry (IHC) and Immuno fluorescence (IF)

For immunohistochemistry, H&E standard procedure was followed, and then ileum sections were incubated with primary antibody for CX3CR1 (1:200; abcam [ab8021] purchased from Saiao, China), CD11a (1:200; biorbyt orb [385445] UK), CD103 (1:200; Absin [118723] purchased from Saiao, China), F4/80 (1:200; Cell signaling technology, [Q61549] US, purchased from Saiao, China), NOD2 (1:200; Novus Biologicals, NB100-524SS), and APC (1:200; Abcam [ab15270], purchased from Saiao, China), for 30 mins, and then stained with diaminobenzidine followed by hematoxylin. Negative controls were incubated only secondary antibody; and showed no significant staining from all cases. Images were captured through Nikon Labophot 2 microscope (ImagePro Plus 3.0, ECLIPSE80i/90i; Nikon, Japan), analyses were performed for expression levels of F4/80, CD11a, CD103, CX3CR1.

Immunofluorescence was performed as already described procedure [38]. Briefly, sections were incubated with primary antibody NOD2 (1:100) and APC (1:100) over night at 4°C, then conjugated secondary antibody (Goat anti mouse Alexa Fluor 594 for NOD2, Donkey anti Rabbit for APC Alexa Fluor 488) respectively at room temperature for 1 hr in the dark and then covered slip with DAPI medium. Images were taken by Nikon Labophot 2 microscope (ImagePro Plus 3.0, ECLIPSE80i/90i; Nikon, Japan).

### 2.4. Tissue preparation for TEM analysis

Ileum (Control and high dose group of 6 months only) was separated from the small intestine and then fixed at room temperature (RT) for 2 hrs. Then, the ileum tissues were washed with PBS, and then post-fixed with osmium tetroxide for 2 hrs at room temperature, followed by 10 min pre-staining in acetate-barbitone. After that dehydrated by graded ethanol, the samples were fixed in Spurr’s resin. Sections were stained with uranyl acetate and lead citrate, and then ultra-structure of dendritic cell migration and bacteria were observed through the JEM-1400 (JEOL Ltd., Tokyo, Japan) TEM.

### 2.5. Bacterial DNA Isolation and 16S rRNA gene Sequencing

During necropsy, mice fecal pellets were collected from the 6 months As exposed mice and then total bacterial DNA was extracted from fecal pellets by using power soil DNA isolation kit (MO BIO Laboratories, Beijing, China). The DNA purity and quantity was estimated at 260/280 nm and 260/230 nm respectively and then sample were preserved at -800C for further analysis. The bacterial 16S rRNA gene (V3 or V6 region) was amplified twice with the suitable thermal cycling conditions and primers (Forward primer, 5’-ACTCCTACGGGAGGCAGCA-3’; reverse primer, 5’-GGACTACHVGGGTWTCTAAT-3’). Finally, the PCR products were analyzed through Quant-iT™ dsDNA HS Reagent and then pooled together. High-throughput sequencing analysis of bacterial rRNA genes was performed on the purified, pooled sample using the Illumina Hiseq 2500 platform (2×250 paired ends) at Biomarker Technologies Corporation, Beijing, China. This method already described in our previous studies (40). (Chiranjeevi et al., 2020)

### 2.6. Separation of epithelial cells from intestine for mRNA extraction

Epithelial cells from ileum were isolated. Briefly, ileum part was separated from the entire small intestine and placed in cold PBS. The ileum tissues were flushed with cold PBS to wash, using 3 ml insulin syringe. Pieces were incubated in 50-100 ml of 0.04% sodium hypochlorate on ice for 15 min to avoid bacterial contaminations. Later intestinal pieces from sodium hypochlorate solution were rinsed in PBS and then incubated in 15 ml conical tube containing solution B for 15 min on ice. Solution B was discarded and then washed with 5ml of PBS or solution B for two times. Then the intestinal pieces were transferred to another conical tube (containing solution B) and then centrifuged at 1,000 rpm (4°C) for 15min. Then, the cells contained pellet was suspended in 1 ml of Trizole for RNA extraction.

### 2.7. Extraction of mRNA from epithelial cells and Real-time-qPCR

Total RNA was isolated from epithelial cells by using Trizole Reagent according to manufacture instructions and then checked their intensity through agarose gel electrophoresis and the RNA quality and quantities were identified through Nanodrop ND-1000 Spectrophotometer. The RNA was converted to cDNA through the reverse transcription PCR according to the manufacturer’s instructions (PrimeScript® RT Master Mix Kit). The expression levels of genes were quantified by qRT-PCR used by SYBER Premix Ex TaqTM II QRT-PCR kit performed through the Mx3000PTM RT-PCR system, with the suitable primers and thermal profile conditions. This experiment has been done in triplicate, and then obtained raw data was analyzed by the 2-ΔΔCt method.

### 2.8. Protein extraction and analysis of cytokines by ELISA

For ELISA analysis, protein content was isolated by adding 2 ml of PBS (containing 0.1% PMSF) to the intestinal epithelial cells and then allowed to incubate for 10 min under ice condition for the disruption and then centrifuged at ¬15000 rpm for 10 min. The resultant protein supernatant thus obtained was stored at -300C for further process. The anti and inflammatory cytokines (TNF-α, IFN-γ, IL-18 and IL-10) were analyzed through ELISA method according to the manufacture instructions (Westang Biotech Co. Ltd, Shanghai, China). Finally, the plates were read by SCO GmbH (Dingelstadt, Germany). The absorbance obtained from plate reader was transformed to cytokine concentrations (pg/ml) and then determined protein concentration by using a standard curve computed on Excel. The sensitivities for all cytokines were between 4 to 6 pg/ml.

## 3. Statistical analysis

All the experimental data has shown by the mean ± SEM and calibrated with software (GraphPad Software Inc., San Diego, USA) of Prism 5 GraphPad. Significant changes between the control and treatment groups were analyzed by one-way ANOVA followed by a Tukey’s Multiple Comparison test. The considered significant statistical value is p < 0.05. Moreover, for the microbiome overlap relation between paired-end (PE) reads, the double-end sequence data was obtained by Hiseq sequencing merged into sequence tags, quality of reads; effect of merge quality controlled and filtered was analyzed by FLASH v1.2.7, Trimmomatic v0.33, and UCHIME v4.2 software respectively. OTU (Operational Taxonomic Unit) analysis UCLUST[2] in QIIME[1] (version 1.8.0) software was used to cluster the tags and obtain OTU at 97% similarity level, and then prepared taxonomic annotation of OTU based on Silva (bacteria) and UNITE (fungi) taxonomic database. In addition, the sequences of OTU with the highest abundance at the taxonomic level were selected with the QIIME software as the representative sequences, and then the multiple sequence alignment was conducted and the phylogenetic tree was constructed. Then the graph was drawn with the Python language tool, Mothur (version v.1.30) software was used to evaluate the sample Alpha diversity index.

## 4. RESULTS

### 4.1. Gut-microbiome alterations induced by As

Figure 1 revealed that abundance of gut bacteria indicated at family level through the 16S rRNA sequencing states, each color indicating a specific bacterial family. Among the bacterium allocating level of phylum, Bacteroidetes (60.0.4%) and Firmicutes (30.10%) were predominant in the gut bacteria of mice, followed by Proteobacteria (8.58%), Deferribacteres (0.56%), Actinobacteria (0.31%), Tenericutes (0.29%), and Saccharibacteria (0.11%). Our results at phylum level are in correlation with earlier studies by Turnbaugh et al. (2006). Significant fold changes (p<0.05) and taxonomical assignment of gut bacterial components were recorded in figure (1E). Impact of As on the alteration of gut microbiome was significant in high dose group (Fig 1G) when compared with the control group. Control and As exposure animals were well differentiated with 41.04% and 31.20% variation, this variation demonstrated through principal component 1 and 2, respectively. Control and As exposure animals clustered in their individual groups, were demonstrated through the PCoA plot, by the analysis of evolutionary diversity UPGMA, as shown in figure 1D. Moreover, control mice divided into two subgroups in the PCoA plot (Fig. 1 G) and UPGMA analysis (Fig. 1D), which could be due to the own differentiation of microbiome profiles, as demonstrated in figure 1A.

**Figure 1.**
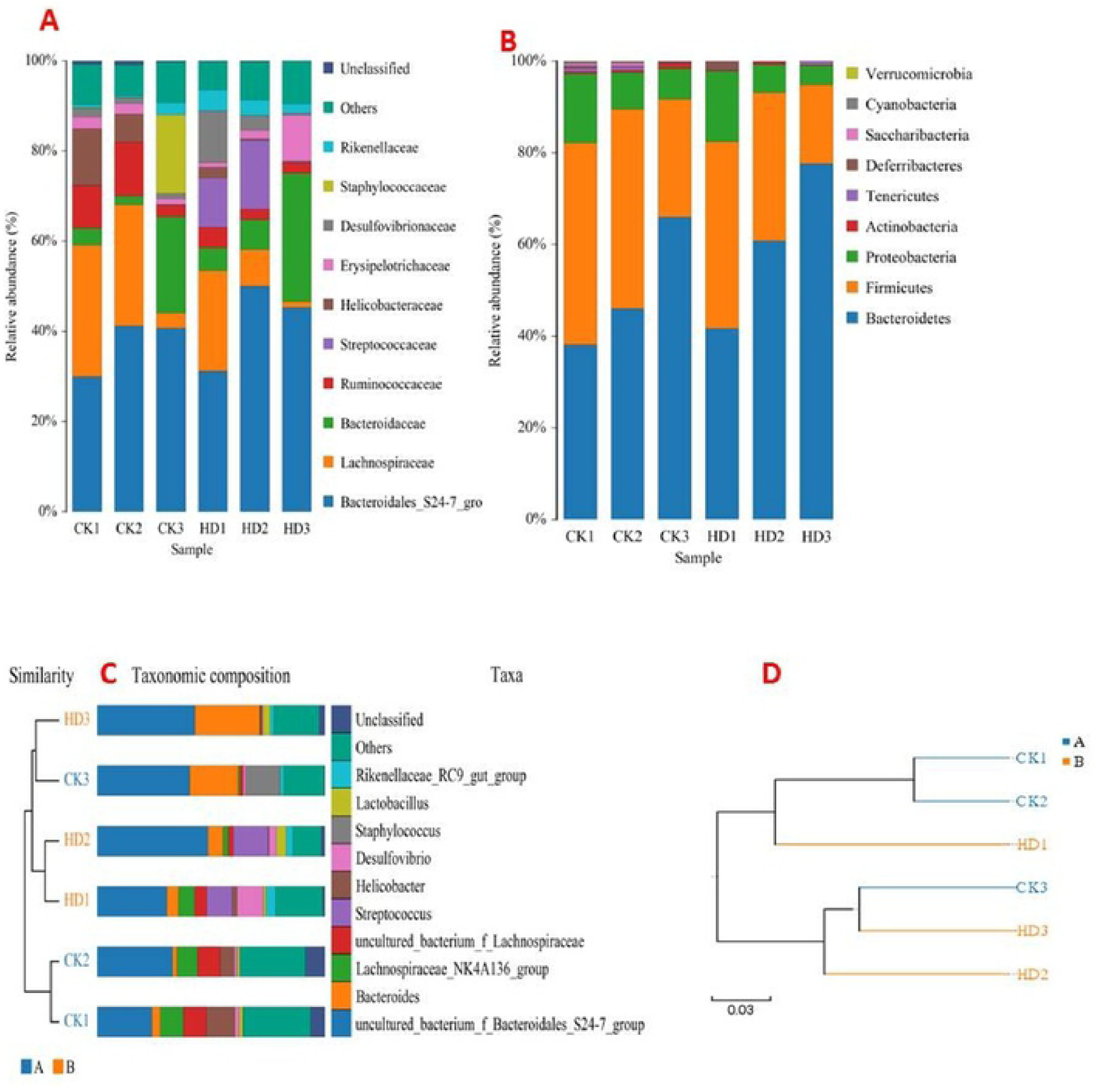
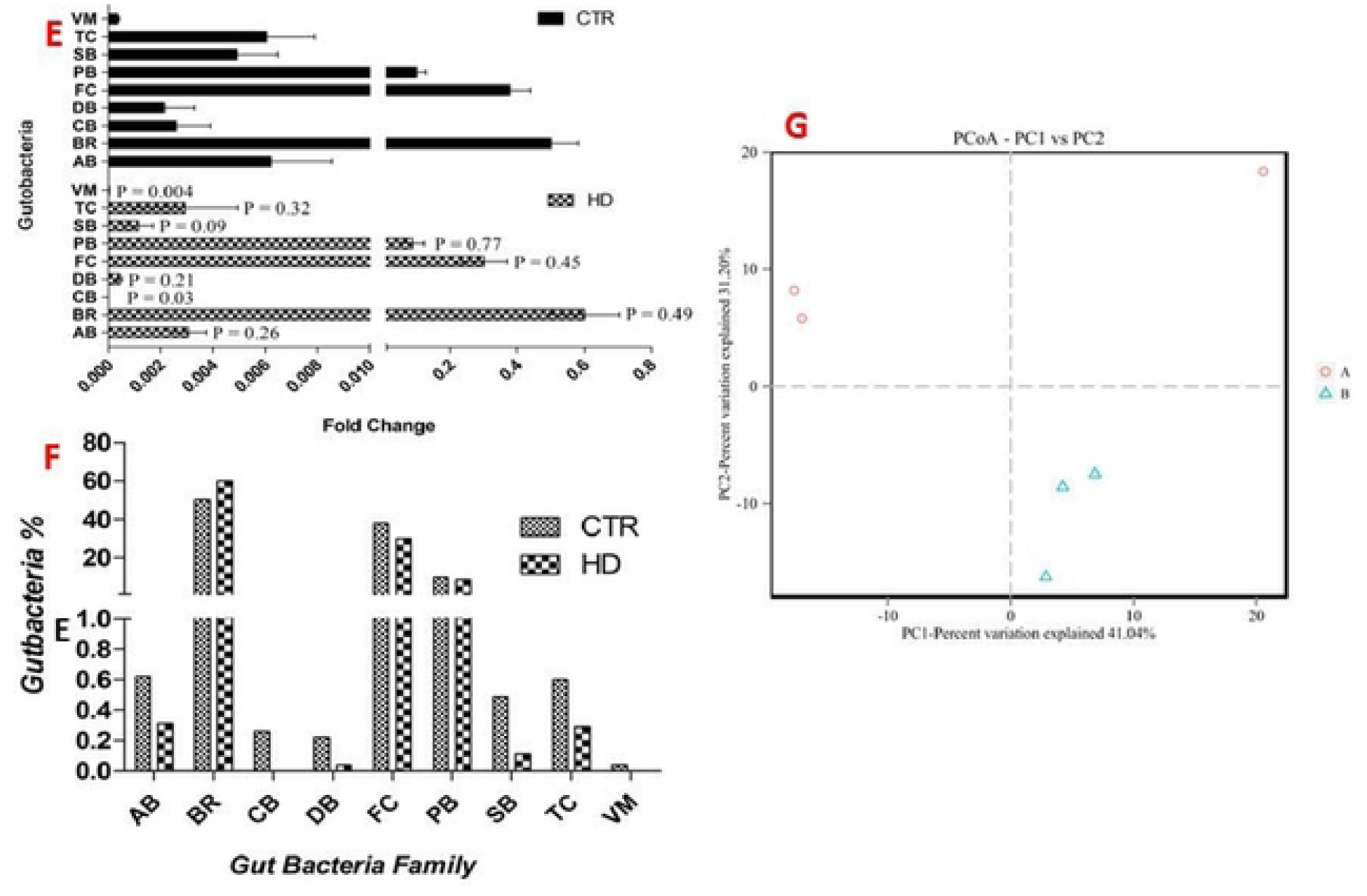
**A)**. Both in control and treated gut microbiome profile evaluated through the l6S rRNA sequencing each color indicating their individual bacterial family (CK= Control; HD= High dose). **B)**. Showing gut bacterial abundance at phylum level. **C)**. The gut bacteria showing taxonomical view. **D)**. the tree both in control and treatment created with 0.03 distance by UPGMA **E)**. Fold changes of gut bacteria showing significant changes of gut bacteria in arsenic treated mice in contrast to controls groups each color indicating specific bacteria at phylum level **F)**. Graph showing the percentage (%) of gut bacteria in both control and As treated animals **G)**. The alteration of gut microbiome in control and arsenic-treated mice varied by PCA circle shape indicating in red color (Control=A) Arsenic high dose, indicating blue color (High dose= B)(Ab = *Actinobacteria*, Br = *Bacterodietes*, Bb = *Cyanobacteria*, Db= *Deferribacteres*, Fc= *Firmicutes*, Pb = *Proteobacteria*, Sb = *Saccharibacteria*, Tc= *Tenericutes*, Vm = *Verrucomicrobia*).

### 4.2. Dendritic cells tried to engulf the pathogenic bacterium

The dendritic cells (DCs) migrate from epithelium to lumen and protect from the pathogenic bacteria across through the epithelial barrier (Nicoletti, 2000). In this study, we have pursued the ultra-structure of intestinal dendritic cells (DCs), macrophages and bacterial mobilization. Here, we reported the dendritic cells ultra-structure and their migration in 6 months As exposed mice intestine. Our results showed that there was no cellular extension and bacterial structure found in control group (Fig. 2A), while ultrastructure of cellular extension (Fig. 2B) trying to engulf the bacterium (Fig. 2C), at epithelial cells into the intestinal lumen was observed in As treated group. Not only the dendritic cell extension and also bacterium crossing the epithelium barrier (Fig. 2D) to penetrate the cell (Fig. 2E) and fine ultra-structure of macrophage migration (Fig. 2F) were observed in As treated groups. We observed fine ultra-structure of cellular organelles in control group, while the ultimate structure of dendritic cell, macrophages, bacterial structures and their mobility in As treated group.

**Figure 2.**
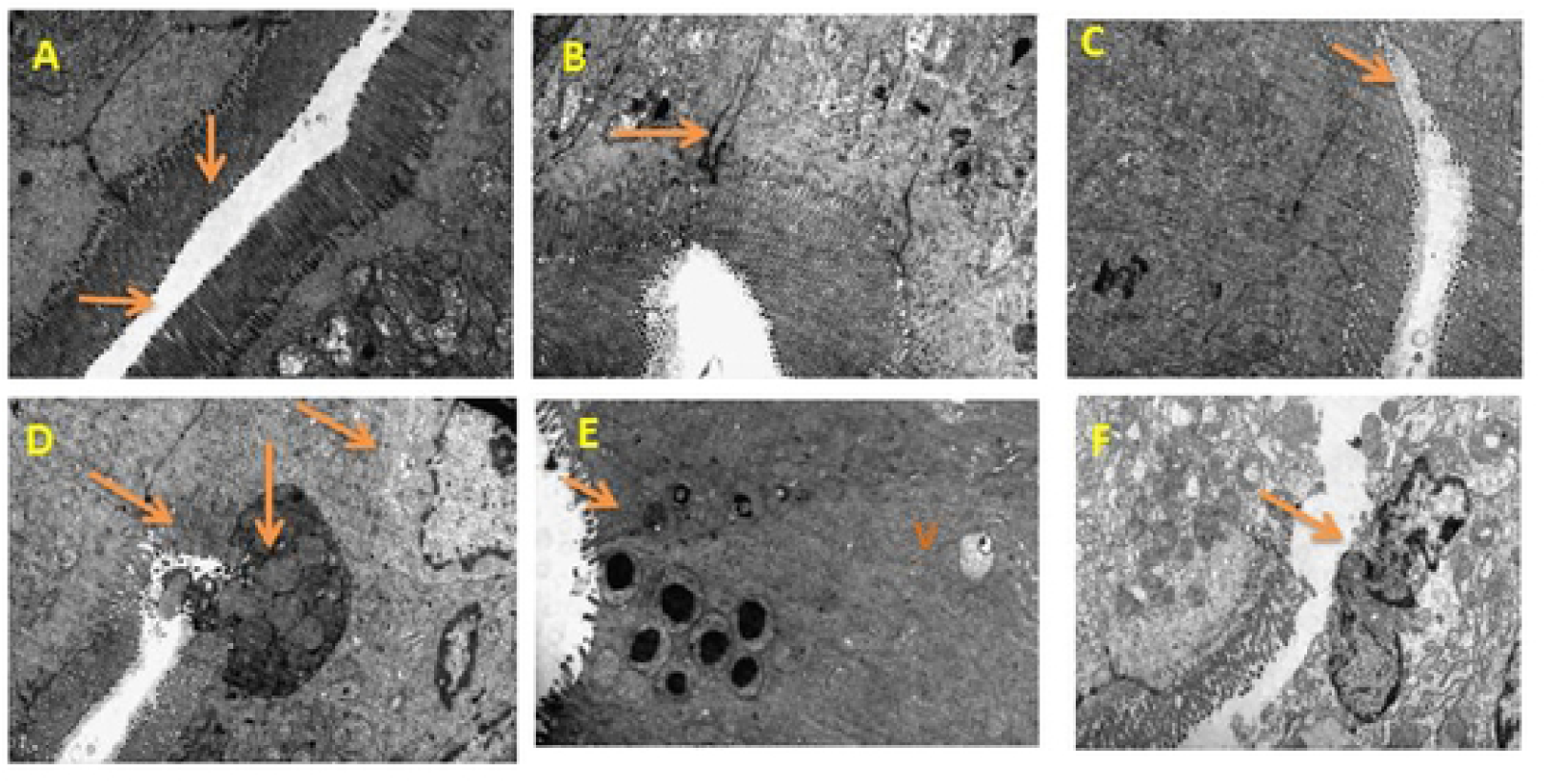
TEM micrograph showing: **A**. Intestinal epithelial cells and villi in control mice. **B**. Dendritic cell migrate to embedded epithelial cells in high dose. **C**. Dendritic cell trying to engulf the Microbial pathogens in high dose mice intestine. **D**. Pathogenic bacteria entered through embedded epithelial cells. **E**. Bacteria encapsulated with vacuole in high dose. **F**. Inflammation formed in epithelial layer of small intestine.

### 4.3. Interrelation between gut micro biome and macrophage (MΦ) F4/80 and CX3CR1

The mRNA expression, protein distribution and intensity pattern were investigated from all four groups through qRT-PCR, immunohistochemistry respectively. In this study, we have studied intestinal immune proteins such as macrophage (MΦ) F4/80 (Fig. 3) and dendritic cell CX3CR1 (Fig. 4). Based on As dose, MΦ and dendritic cells were expressed, however the mRNA levels of F4/80 (Fig. 3G, N) and CX3CR1 (Fig. 4F, M) were significantly increased, moreover the protein intensity F4/80 (Fig. 3A, B, C, D, E, F, H, I, J, K, L, M) and CX3CR1(Fig. 4A, B, C, D, E, G, H, I, J, K, L, N) were significantly distributed both in 3 and 6 months exposure period. But there was no significant differences of F4/80 protein intensity were found in low dose group of both 3 and 6 months exposure mice, while during there was no significant different in 3 months low dose group of CX3CR1 protein intensity but not in 6 months. These differences were explains some animals may addicted to the arsenic due to the prolonged exposure.

**Figure 3.**
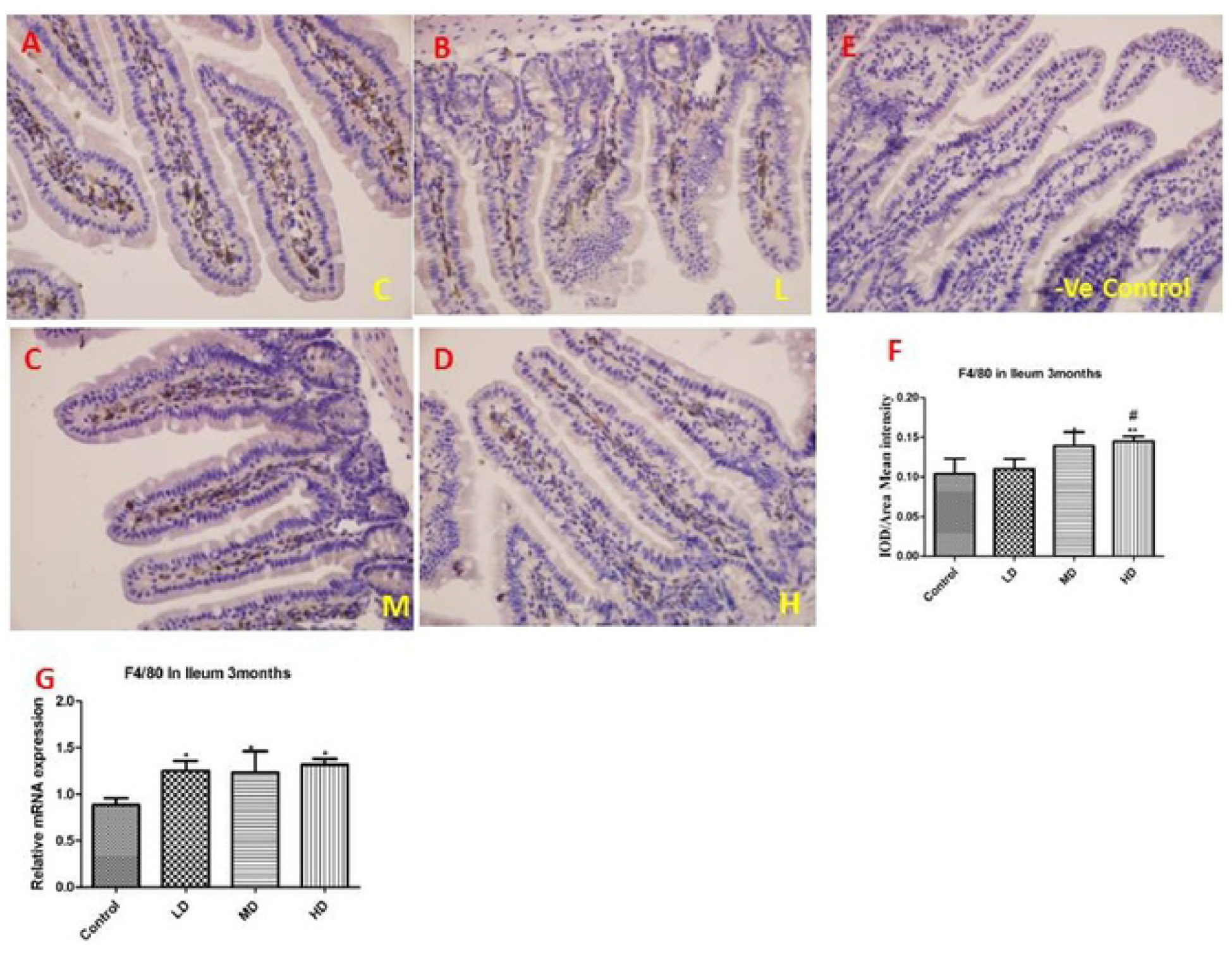
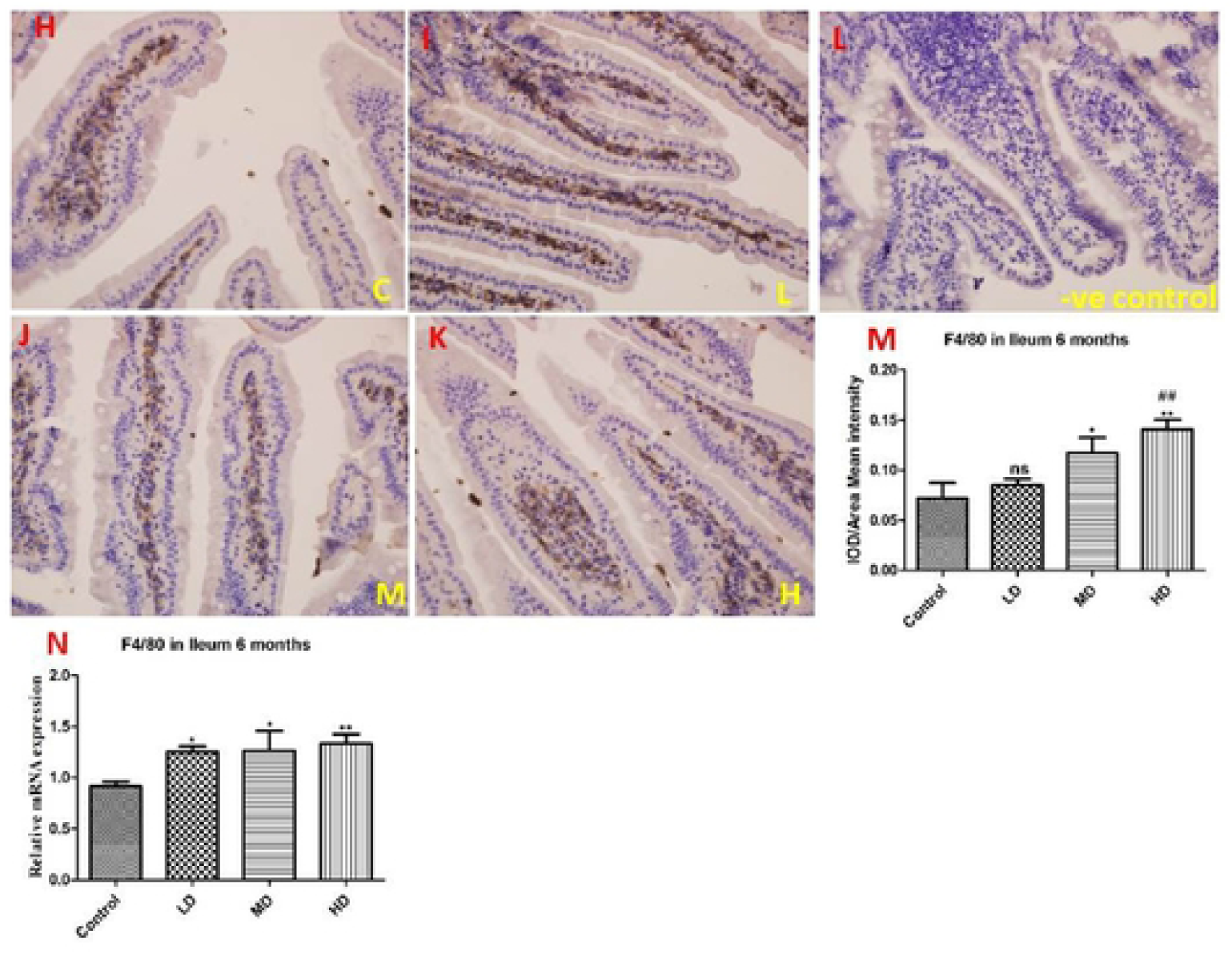
Immuno histochemistry revealed F4/80 intensity, location and expression in 3months Ileum **[A** indicates Control, **B** indicates Low dose, **C** indicates Medium dose, **D** indicates High dose, **E** indicates Negative control, **F** indicates F4/80 protein intensity, **G** indicates Relative mRNA expression level of F4/80], arsenic treated group (LD, MD and HD) compared with control group. In 6months Ileum **(H-**Control; **I**-Low dose group; **J** -Medium dose group; **K**-High dose group; **L-**Negative control; **M**-F4/80 protein intensity; **N**-Re lative mRNA expression level of F4/80].

**Figure 4.**
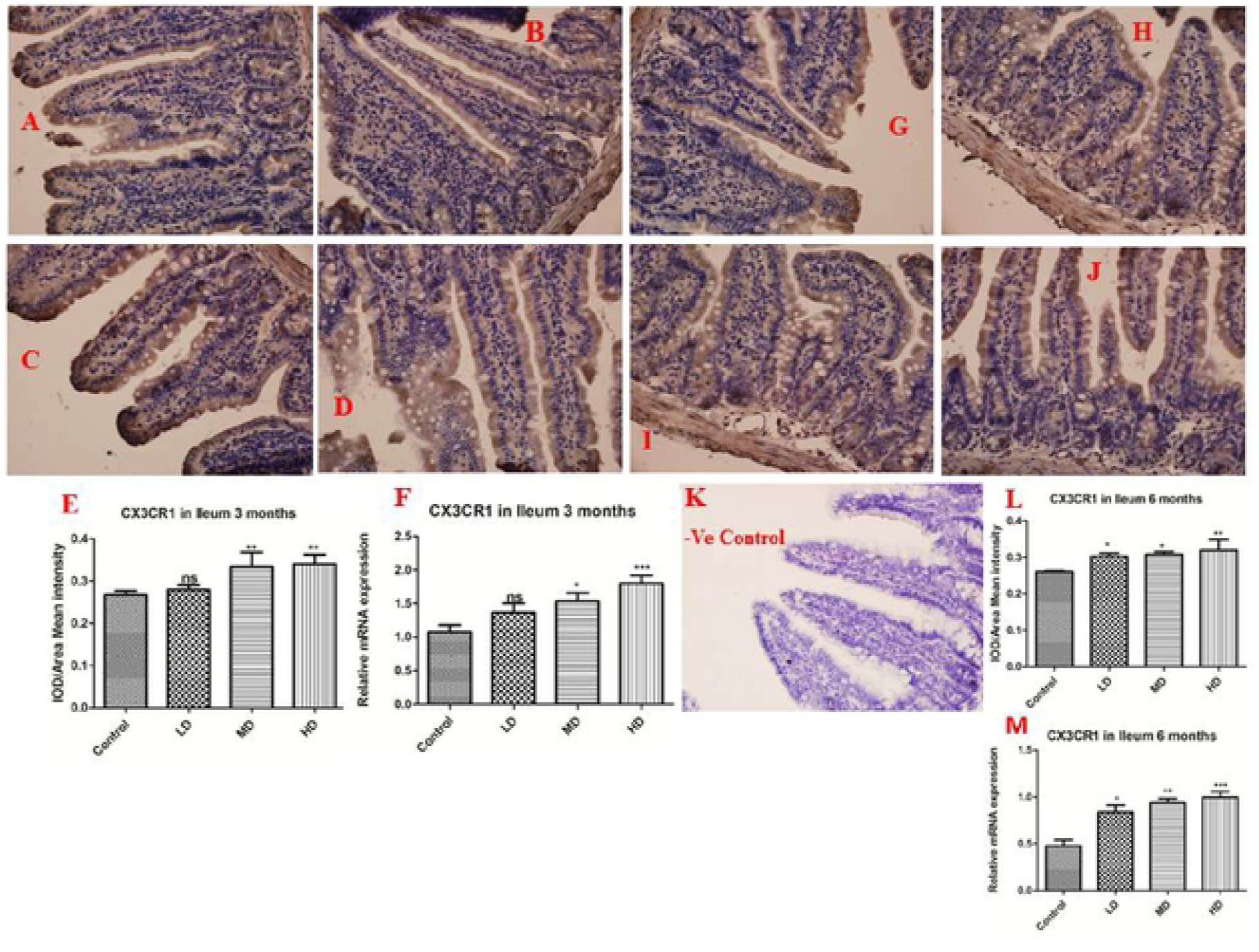
In 3 months Ileum [**A**-Control, **B**-Low dose group, **C**-Medium dose group, **D**-High dose group, **E**-CX3CRI protein intensity bar graph], **F)**. Impact of arsenic on relative mRNA level of CX3CRI in Intestinal epithelium of Ileum 3 months. The intensity of protein were estimated in arsenic treated groups (LD, MD and HD) and compared with control group. **In 6 months Ileum [G-**Control; H-Low dose group; I-Medium dose group; J -High dose group; L-CX3CRI protein intensity bar graph, **K-**Negative control], **M**. Effects of arsenic on relative mRNA level of CX3CRI in Intestinal epithelium of Ileum 6 months. Treated groups (LD, MD and HD) and compared with control group.

### 4.4. The inflammatory cytokines secreted by the altered dendritic cells

Influence of As on the levels of anti-inflammatory and inflammatory cytokines (IFN-γ, IL-18, TNF-α and IL-10) were quantified through the qRT-PCR (Fig.5A, B, C, D, E, F, G, H, I, J, K, L, M, N, O, P). Our results revealed that IFN-γ, IL-18 and TNF-α were significantly increased (Fig. 5), with a concomitant decrease in IL-10 (Fig-5I, J, K, L) in both 3 and 6 months exposure animals. But, no significant differences were identified in case of IFN-γ protein in low dose of 3 and 6 months (Fig. 5C, D), TNF-α mRNA in low dose group of 3 and 6 months (Fig.5E, F); IL-18 mRNA level in both in 3 and 6 months low dose exposed groups (Fig. 5M, N) and IL-10 in 3 and 6 months low dose exposed groups (Fig.5I, J, K). Interestingly, dose- and time-dependent effect of As was observed, i.e., among the three different As doses, high dose showed significant effect when compared with other two doses and moreover, among the two exposure periods studied, the mRNA expression levels were found to be higher in the 6 months age group (Fig. 5) compared with the 3 months age group.

**Figure 5.**
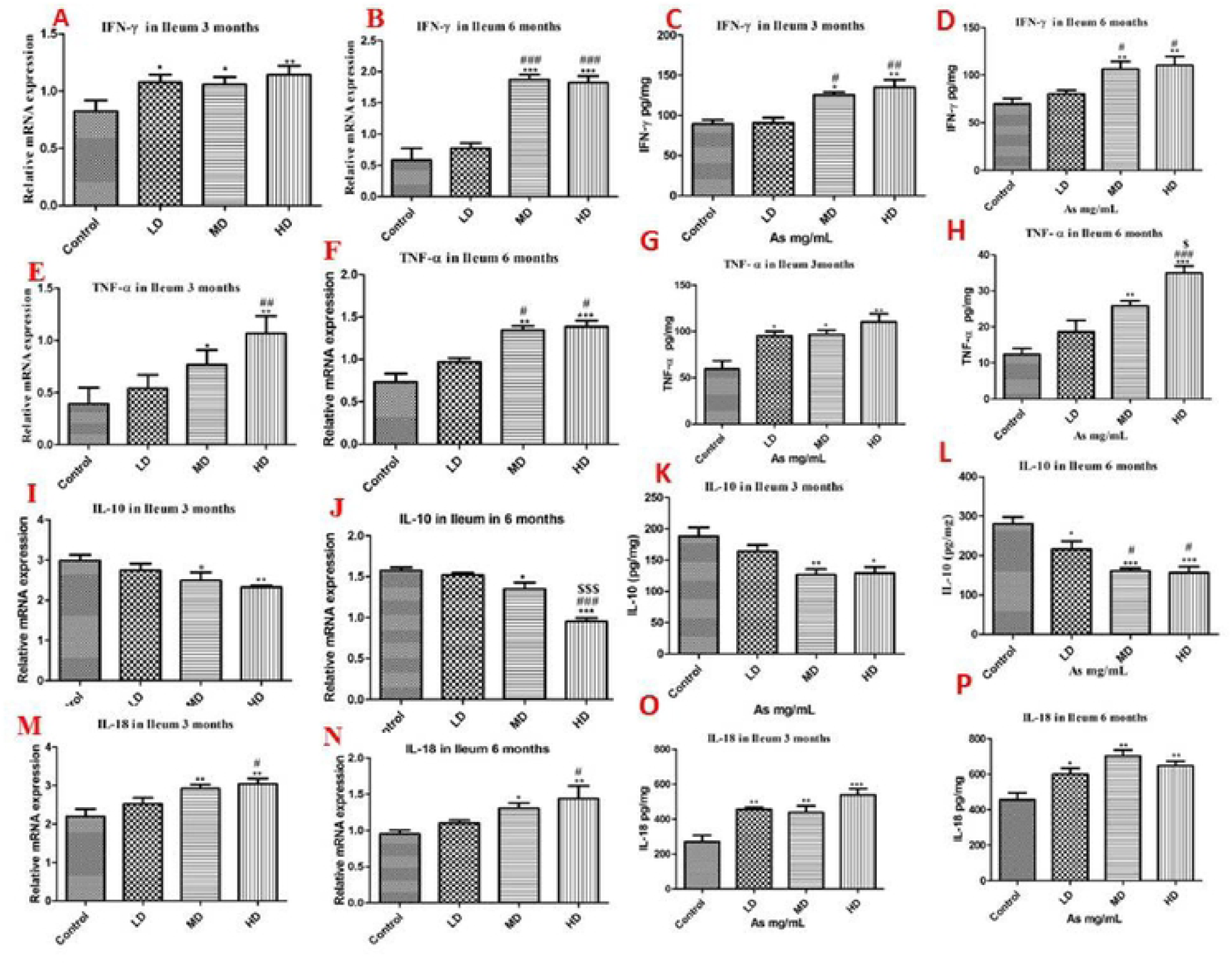
Effects of As on relative mRNA expression levels of Intestinal epithelial cells of Ileum in 3 months and 6 months As exposure and Control groups. **A, B** bar graphs show relative mRNA expression levels **C, D** indicates protein levels of IFNγ in 3 and 6 months As exposure male mice respectively. **E, F**. graph show relative mRNA expression levels **G, H** indicates protein levels of *TNF-α* in 3 months and 6 months As exposure male mice respectively. **I, J**. Graph show relative mRNA expression levels **K, L** indicates protein levels of IL-10 in 3 and 6 months As exposure male mice respectively. **M**, **N**. Bar-graph show relative mRNA expression levels **0**, **P** bar-graph indica tes protein levels of IL-18 in 3 and 6 months As exposure male mice respectively. Data represent the mean±SEM (n=6). Asterisk (*) indicates significant difference compared to the control group (*p < 0.05; **p<0.01), Hash (#) indicates significant difference compared to the low dose group (#p < 0.05; ##p<0.01) and dollar($) indicates significant difference compared to the medium dose group ($p < 0.05; $$p<O.O!) and lack of symbol on any of the bar, except control group, represents non-significant with their respective compared group. LO: low dose, MD medium dose, HD: high dose.

## 5. DISCUSSION

In general, humans are being exposed to very low doses of As, hence to maintain a rationale with environmental As levels, in this study, we have exposed mice with 0, 0.15, 1.5, 15 ppm As_2_O_3_ for 3 and 6 months to evaluate the impact of As on gut microbiome and immune system. The present study has been revealed that impact of As exposure on the gut microbiota, immune system and its mechanism associated with inflammation in small intestine of Ileum. Our data clearly indicated that prolonged exposure of As changed immune system and regulate inflammation through the alteration of gut microbial compositions. In previous studies we provided that arsenic deregulates NOD2 (Nolan Maier et al., 2014) and altered gut microbiome leads to change the immune system and influence the colon cancer marker in small intestine of Jejunum (Lu et al., 2014), Our results provided mechanistic visions regarding alteration of the gut microbiome to disrupt immune system which acts as a novel mechanism of environmental factors-induced human diseases like cancer.

The accumulating evidence suggests that As exposure is linked to altered gut microbiota in children (Dong et al., 2017). Exposure to As significantly depleted alpha diversity in the gut microbiota with a decrease in the bacterial population (Liang Chi et al., 2017). These alterations in the bacterial population could significantly dysregulate the normal gut microbiome functions (Kashyap et al., 2013). Recent studies revealed that repeated doses during early development or adulthood altered gut microbiome linked with immune disruption (Gokulan et al., 2018). Moreover, several other recent studies reported that small intestinal and cecal microbiome alterations (Viaud et al., 2013; Mireia et al., 2018), while the fecal microbiota altered significantly in our study. In consistent with these views, we reported that high As dose altered gut microbiome, while, low dose As doesn’t.

The microbiome is essential to organize intestinal homeostasis. Here we evaluated gut microbiome alterations influenced by the arsenic through high-throughput 16S rRNA gene sequencing. In this study our results clearly showed that chronic exposure to arsenic not only altered gut microbial composition and also changed the intestinal homeostasis, these changes strongly associate with the cytokines modifications. But previous studies provided that arsenic showed impact on gut microbiome alterations and its perturbed gut microbiome strongly associate with changes of many microbial floras ((Lu et al., 2014). Recent studies reviewed that microbiota plays a vital role in the initiation, training and function of the host immune system (Belkaid et al., 2014). The germ-free (GF) animals or any other models organism were not exposed to any pathogenic microbes and thus have a dominant innate and adaptive immune system (Smith et al., 2007).

The gut microbiome regulates immunomodulatory functions through their interactions with TLRs expressed on the surfaces of epithelial cells and DCs (Abreu et al., 2010) and different bacteria stimulate different and distinct TLRs on host cells (Vanderpool et al., 2018). The dysbiosis of microbiota regulates γδT17 cells by the activation of CD103+ leading to drive total monoclonal expansion (Fleming Chris et al,m 2017). Here, we proved that the dysbiosis of microbiota activated DCs which stimulates the inflammatory cytokines production while depleted anti-inflammatory cytokines. In this study, exposure to As revealed that dysbiosis of microbiota activated macrophages (F4/80), and DCs (CX3CR1) to produce inflammatory initiators (IFN-γ, TNF-α, and IL-18) and anti-inflammatory cytokine (IL-10). The imbalance in immune system leads to the secretion of inflammatory cytokines. Some of the other studies reported that increased production of interleukin IL-12 and IL-8 and TNF-α by DCs was significantly associated with survival (Vivek Subbiah et al., 2018). Fernanda et al., (2009) reported that TNF-α, and IL-12 were involved in the TLR4/Smteg interaction of MyD88 signaling pathway due to the activation of DCs by Smteg (Durães et al., 2009). Earlier studies showed that TNF-α, largely secreted by Ly6c+ CD11b+ dendritic cells (DCs), plays a vital role in promoting IL-17A from CD4+ T cells and associated to induce airway neutrophilia (Feia et al., 2011). There has been controversy with respect to dendritic cells such as macrophages, capable to produce IFN-γ (De Saint-Vis et al., 1998). Previous reports illustrated that CD8a2 and CD8a1 splenic dendritic cells produced IFN-g in response to IL-12p70 (Ohteki et al., 1999), an effect that was inhibited by the accumulation of IL-4 or IL-18 (Fukao et al., 2000). Some of the studies reviewed that, the pro-inflammatory cytokines (Th1 and Th2) are produced by the cytokines and chemokine signaling pathway (Pierre Miossec et al., 2008). Such kind of inflammation initiators create cellular modifications that can promote to chronicity. The present report determined the regulatory mechanism of inflammatory interleukin and anti-inflammatory cytokine expressions by dendritic cells. IL-10, IL-18, IFN-γ and TNF-α transcripts detected by real-time PCR were promptly up-regulated by dendritic cells CX3CR1 and macrophage F4/80 stimulation. Furthermore, IL-10 was depleted by the stimulation of dendritic cells and macrophage while regulated IL-18 IFN-γ and TNF-α. These results were correlated with previous studies by Xavier and Podolsky (Xavier et al., 2007). The gut inflammatory pathway regulates different types of cancers like colon and rectal cancer (Neuman 2007; Terzic et al., 2010). Moreover, addition of proinflammatory cytokines helps in the formation of inflammatory state in the colon and to promote colon and rectal cancer (Sanchez-Munoz et al., 2008).

Kristina et al., (2012) described the role of anti-inflammatory and inflammatory cytokines in colon and rectal cancers. Furthermore, recent reports evidenced that activation of β-catenin signaling in effector T cells and/or Trigs is causatively linked with the marking of proinflammatory properties and the upgrade of colon cancer (Shilpa Keerthivasan et al., 2014). Our results showed that As exposure altered gut microbiome composition and strongly associated with changed related microbial flora, regulate to activate macrophages and dendritic cells. The irregular function of intestinal homeostasis secreted the inflammatory cytokines in male mice and which provides mechanistic information to determine how chronic exposure of As influence on the gut microbiome, immune system and regulate inflammation.

## 6. Conclusions

In conclusion, we have proved that chronic exposure to arsenic strongly associated with inflammation. However, arsenic influenced the inflammation through the alteration of gutmicrobiome and changes of intestinal immune system. The dysbiosis of gut microbiota leads to significantly increased both mRNA and protein levels of macrophages (F4/80) and dendritic cells (CX3CR1) while decreasing the anti-inflammatory cytokines (IL-10) and increased level of inflammatory cytokines (IFN-γ, TNF-α, and IL-18). Altered immune system and regulate the intestine inflammation, providing novel platform to determine how chronic exposure to As leads to various diseases. In previous studies we have reported on the alteration of gut fungus and immune system in the intestine part of Jejunum (Chiranjeevi Tikka et al., 2020).

## Notes

The authors declare that there are no conflicts of interest

